# Bio-hybrid soft robots with self-stimulating skeletons

**DOI:** 10.1101/2020.09.16.299719

**Authors:** Maria Guix, Rafael Mestre, Tania Patiño, Marco De Corato, Giulia Zarpellon, Samuel Sánchez

## Abstract

Bioinspired hybrid soft robots combining living actuation and synthetic components are an emerging field in the development of advanced actuators and other robotic platforms (i.e. swimmers, crawlers, walkers). The integration of biological components offers unique properties (e.g. adaptability, response to external stimuli) that artificial materials cannot replicate with accuracy, being skeletal and cardiac muscle cells the preferred candidates for providing contractile actuation. Here, we present a skeletal-muscle-based swimming biobot with a 3D-printed serpentine spring skeleton that provides mechanical integrity and self-stimulation during the cell maturation process. The restoring force inherent to the spring system allows a dynamic skeleton compliance upon spontaneous muscle contraction, leading to a novel cyclic mechanical stimulation process that improves the muscle force output without external stimuli. Optimization of the 3D-printed skeletons is carried out by studying the geometrical stiffnesses of different designs *via* finite element analysis. Upon electrical actuation of the muscle tissue, two types of motion mechanisms are experimentally observed: i) directional swimming when the biobot is at the liquid-air interface and ii) coasting motion when it is near the bottom surface. The integrated compliant skeleton provides both the mechanical self-stimulation and the required asymmetry for directional motion, displaying its maximum velocity at 5 Hz (800 micrometer second^−1^, 3 body length second^−1^). This skeletal muscle-based bio-hybrid swimmer attains speeds comparable to cardiac-based bio-hybrid robots and outperforms other muscle-based swimmers. The integration of serpentine-like structures in hybrid robotic systems allows self-stimulation processes that could lead to higher force outputs in current and future biomimetic robotic platforms.

## Introduction

Biological systems have evolved throughout millennia to develop sophisticated mechanisms of self-organization ^[1]^, actuation ^[2]^, self-healing ^[3]^, and sensing ^[4]^. The robotics field aims to mimic and incorporate these complex behaviours ^[5–8]^. In particular, biomimetic soft robotics research seeks to make programmable flexible and compliant architectures able to actuate in a gentle manner, posing as main challenges the achievement of certain mechanical properties (i.e. compliance, flexibility) and sensing capabilities to comply with the overall safety for human interaction, as the rigidity and stiffness of conventional materials limit their applications into healthcare or biomedical disciplines. Recent developments in material science enabled the fabrication of biomimetic soft robots that are able to perform some simple types of actuation ^[9]^, including crawling ^[10,11]^ or grasping ^[12]^, but they are still far from the degree of complexity and sophistication of their biological counterparts.

One of the most investigated applications in soft robotics is the development of artificial muscles that can mimic the performance of native muscle tissue ^[6]^. Muscle tissue is inherently complex, as it is simultaneously strong and fast, enabling a wide variety of movements through an efficient self-organization of its fiber bundles. Materials of synthetic origin still lack the ability to fully replicate these properties ^[13]^, although recent advances based on pneumatic or dielectric actuators ^[14,15]^, the incorporation of nanomaterials ^[16,17]^ or exploring alternative helical fiber configurations have been reported ^[18]^. Even more, other features from biological tissues, such as self-healing, energy efficiency, power-to-weight ratio, adaptability or bio-sensing, are strongly desired but difficult to achieve with artificial soft materials ^[19]^. Bio-hybrid robotics is born at this point as a synergistic strategy to integrate the best characteristics of biological entities and artificial materials into more efficient and complex systems that can overcome the difficulties that current soft robots face ^[20,21]^. Therefore, the design of bio-hybrid robots relies on the combination of living cells/tissues with a synthetic material, and such living entities can range from individual motile cells at the microscale (i.e. sperm ^[22]^, bacteria ^[23]^), to engineered single cells (i.e. muscle cells), multicellular systems (i.e. co-culture of neurons and muscles ^[24]^, or passive and contractile tissue from *Xenopus laevis* ^[25]^).

Initial examples of bio-hybrid actuators comprised the integration of cardiac tissue were based on spherical heart pumps, 2D cultures that could deform PDMS cantilevers for real-time force monitoring or crawling robots with different amounts of legs that could move due to friction differences ^[26–31]^. Recent advancement in both 3D printing ^[32]^ and tissue engineering ^[33]^ allowed the creation of on-demand functional biological structures, paving the path to their integration in smart soft robotic platforms. Regarding the use of cardiac cells to power bio-robotic devices, most of the fabricated biobots have been based on thin film structures, normally PDMS, seeded with cardiac cells ^[34]^, promoting a two-dimensional layer configuration have been extensively explored for fish-like swimming purposes ^[35,36]^, mimicking the motion of their biological counterparts. Some examples include a swimming bio-hybrid soft robotic sting-ray based on optogenetically modified cardiac cells ^[37]^ or a cardiac-based medusoid that mimics the thrusting mechanism of a jellyfish ^[38]^. Spontaneous contractions present in cardiac cells, whose response can be stimulated and synchronized at a certain frequency, cannot be stopped, limiting the possibility of finer control over the actuation of cardiomyocyte-based bio-hybrid robots. In contrast, skeletal muscle tissue possesses a wider range of adaptability and controllability, since these cells adopt a three-dimensional structure that can be accommodated to different substrates and their contractions are induced on-demand by means of an external electrical or optical stimuli ^[39,40]^.

While cardiac muscle-based living robots are mostly swimmers, in the case of skeletal-muscle-based biobots most examples rely on actuators or walkers ^[41,42]^. Some of the reported tethered skeletal muscle bio-hybrid actuators owed its motion to the deflection of cantilevers by single myotubes, full tissue ^[43–47]^ or as grippers ^[41,48]^. However, for untethered bio-hybrid robots, crawling has been the main described motion mechanism ^[49]^. For example, a 3D-printed biobot composed of two legs joined by a beam structure demonstrated walking motion on the bottom of a Petri dish by differences in friction between two asymmetric legs when the muscle-based actuator was stimulated ^[42]^. Later, the same bio-robotic device was light-controlled remotely by optogenetically-modified skeletal muscle cells which contract upon blue light stimulation ^[40]^. In addition, this device demonstrated self-healing ^[50]^, adaptability ^[40]^, integration of motor neurons for advanced stimulation ^[51]^, long-time preservation ^[52,53]^, scalability ^[54]^ or their integration with micro-electrodes ^[55]^. The integration of neuronal and skeletal muscle tissue in one single biobot has been of great interest, as it resembles the structure of native muscle to obtain improved controllability of the bio-robotic systems ^[24,56]^. In this regard, a bio-hybrid swimmer with functional neuro-muscular junction that swims with time-irreversible flagellar dynamics has recently been reported ^[57]^. This biobot configuration represents the first swimmer based on optogenetically modified skeletal muscle tissue, whose actuation relies on external light stimuli and it presents slow swimming speeds (0.92 µm/s) when moving at low Reynolds number regime. Although several aspects concerning skeletal muscle based biobots controllability and scalability had been explored, their motion mechanism mainly relies on walking or crawling, being key to find alternative efficient motion mechanisms such as swimming to develop faster and more versatile robotic platforms.

Here, we report a skeletal muscle-based swimming biobot with enhanced force performance and directional motion. The integration of a serpentine spring skeleton into the biobot platform allows proper mechanical integrity of the whole system, as well as mechanical self-stimulation due to the spring restoring force when spontaneous contractions take place during the cell maturation process. Such self-training event leads to enhanced actuation and larger contraction force in the biobot performance. This biobot is based on a three-dimensional muscle structure instead of thin films as used in previous cardiomyocyte-based bio-hybrid swimmers ^[37,38]^, offering a wider range of customization and actuation modes. Furthermore, the 3D serpentine spring structure that forms the compliant skeleton has been designed to present asymmetric stiffness throughout its structure, leading to a well-controlled bending of the biobot that permits two different motion modalities: (i) swimming when located at the air-liquid interface, and (ii) coasting when it is placed near the bottom surface. Corresponding motion mechanisms were established by motion tracking and simulations of the locomotion hydrodynamics, revealing that the asymmetry in geometrical stiffness provides directional motion for the swimming case. In fact, our biobot design is the fastest skeletal muscle-based swimming bio-hybrid robot up to date by several orders of magnitude (791x, **Table 1**) and its velocity compares favourably with the bio-robotic systems based on cardiac cells. A novel integration of a serpentine spring on a bio-robotic platform serves both as a compliant skeleton useful for mechanical self-stimulation purposes and to provide asymmetry to the system. The versatility of 3D printing techniques, allowing for rapid and cost-efficient fabrication, added to the properties of this simple yet efficient flexible serpentine spring system demonstrates its potential for a better differentiation and performance of contractile cell-actuated robotic devices, inspiring future bio-hybrid robotic designs with higher efficiency and able to achieve more complex motion patterns.

**Table 1:**
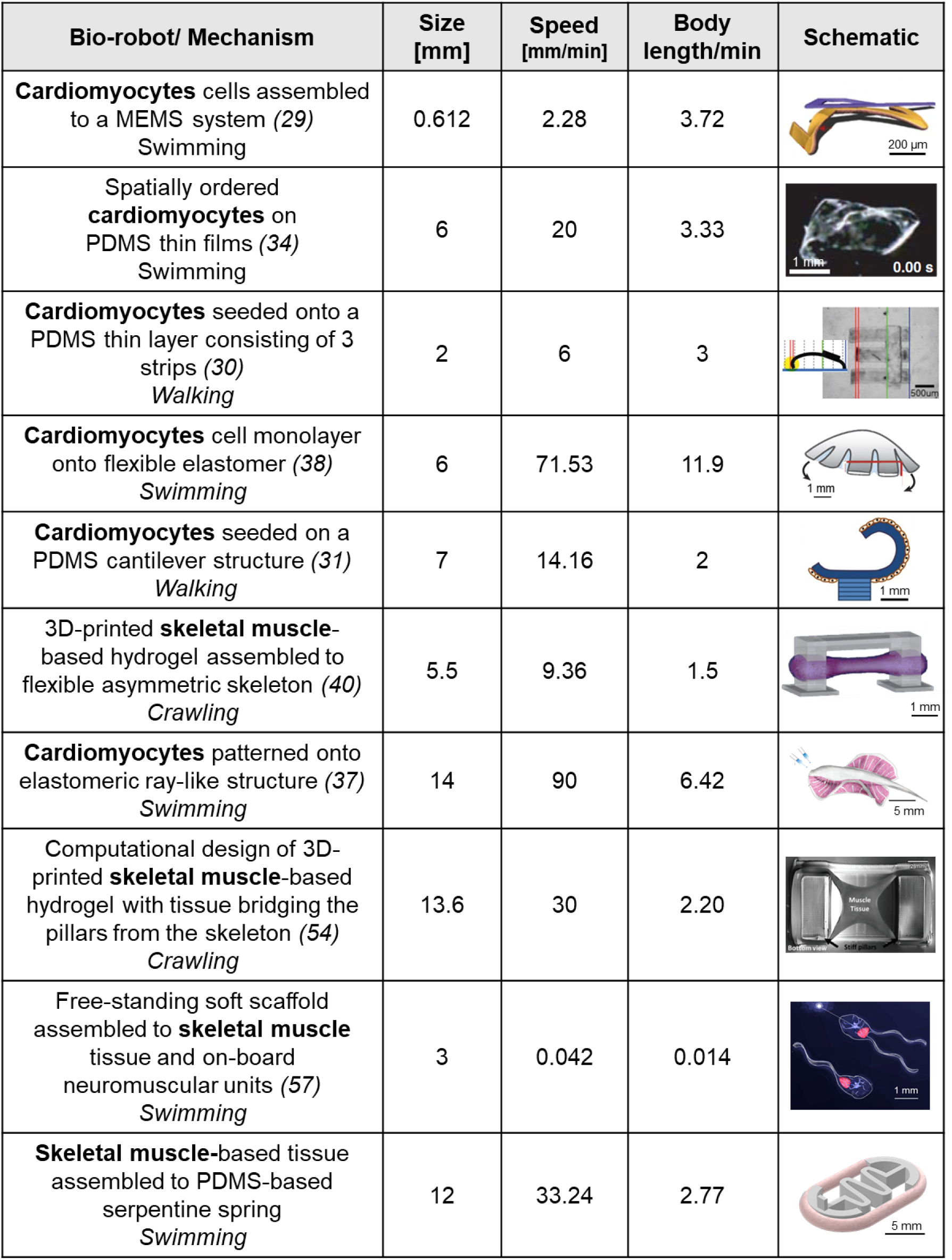
Chronological summary of bio-hybrid robotic systems development based on cardiac and muscular cells, indicating their size, real and relative speed (body length/second), and the corresponding schematic.

| Bio-robot/ Mechanism | Size [mm] | Speed [mm/min] | Body length/min | Schematic |
| --- | --- | --- | --- | --- |
| **Cardiomyocytes** cells assembled to a MEMS system (29)<br>Swimming | 0.612 | 2.28 | 3.72 |  |
| Spatially ordered **cardiomyocytes** on PDMS thin films (34)<br>Swimming | 6 | 20 | 3.33 |  |
| **Cardiomyocytes** seeded onto a PDMS thin layer consisting of 3 strips (30)<br>Walking | 2 | 6 | 3 |  |
| **Cardiomyocytes** cell monolayer onto flexible elastomer (38)<br>Swimming | 6 | 71.53 | 11.9 |  |
| **Cardiomyocytes** seeded on a PDMS cantilever structure (31)<br>Walking | 7 | 14.16 | 2 |  |
| 3D-printed **skeletal muscle**-based hydrogel assembled to flexible asymmetric skeleton (40)<br>Crawling | 5.5 | 9.36 | 1.5 |  |
| **Cardiomyocytes** patterned onto elastomeric ray-like structure (37)<br>Swimming | 14 | 90 | 6.42 |  |
| Computational design of 3D-printed **skeletal muscle**-based hydrogel with tissue bridging the pillars from the skeleton (54)<br>Crawling | 13.6 | 30 | 2.20 |  |
| Free-standing soft scaffold assembled to **skeletal muscle** tissue and on-board neuromuscular units (57)<br>Swimming | 3 | 0.042 | 0.014 |  |
| **Skeletal muscle**-based tissue assembled to PDMS-based serpentine spring<br>Swimming | 12 | 33.24 | 2.77 |  |

## Results

Skeletal muscle-based biobots with an integrated spring-like skeleton were constructed by assembling a cell-laden circular hydrogel around the compliant skeleton (**Fig. 1A**). The skeleton was composed of PDMS and created by extrusion-based 3D printing (**Fig. 1B**). The use of the 3D printing technique provided high versatility and fast prototyping, allowing the design and optimization of different configurations of the artificial skeleton. The main configurations considered were a symmetric (**Fig. 1C1**) and an asymmetric (**Fig. 1C2**) design, the latter including a small post bulging out of one of the sides. This element induced an asymmetric compaction of the skeleton of the biobot, necessary to achieve directional motion. The overall stiffness of the skeleton was tuned by varying the curing agent-to-base ratio of PDMS, but also an extra level of optimization was assessed by modifying its geometrical stiffness. Therefore, different serpentine designs in the central part of the biobot skeleton were considered and further evaluated (**Fig. 1C3**). A rounded notch was included at both ends of the skeleton, being carefully designed and 3D-printed (**Fig. 1C4**) to hold the tissue ring in place and avoid its release during the maturation process or their motion evaluation. Finally, further optimization could be achieved by including different numbers of coils in the design to create biobots with different properties and sizes (**Fig. 1C5**). While the shorter skeleton design did not provide enough flexibility to the system, the design with higher number of coils led to a non-stable assembly of the scaffold to the skeleton due to its large bending at earlier stages. The optimal two-coil configuration, however, allowed an optimal balance between the restoring force and the biobot stability during the maturation process and later motion studies.

**Fig. 1.**
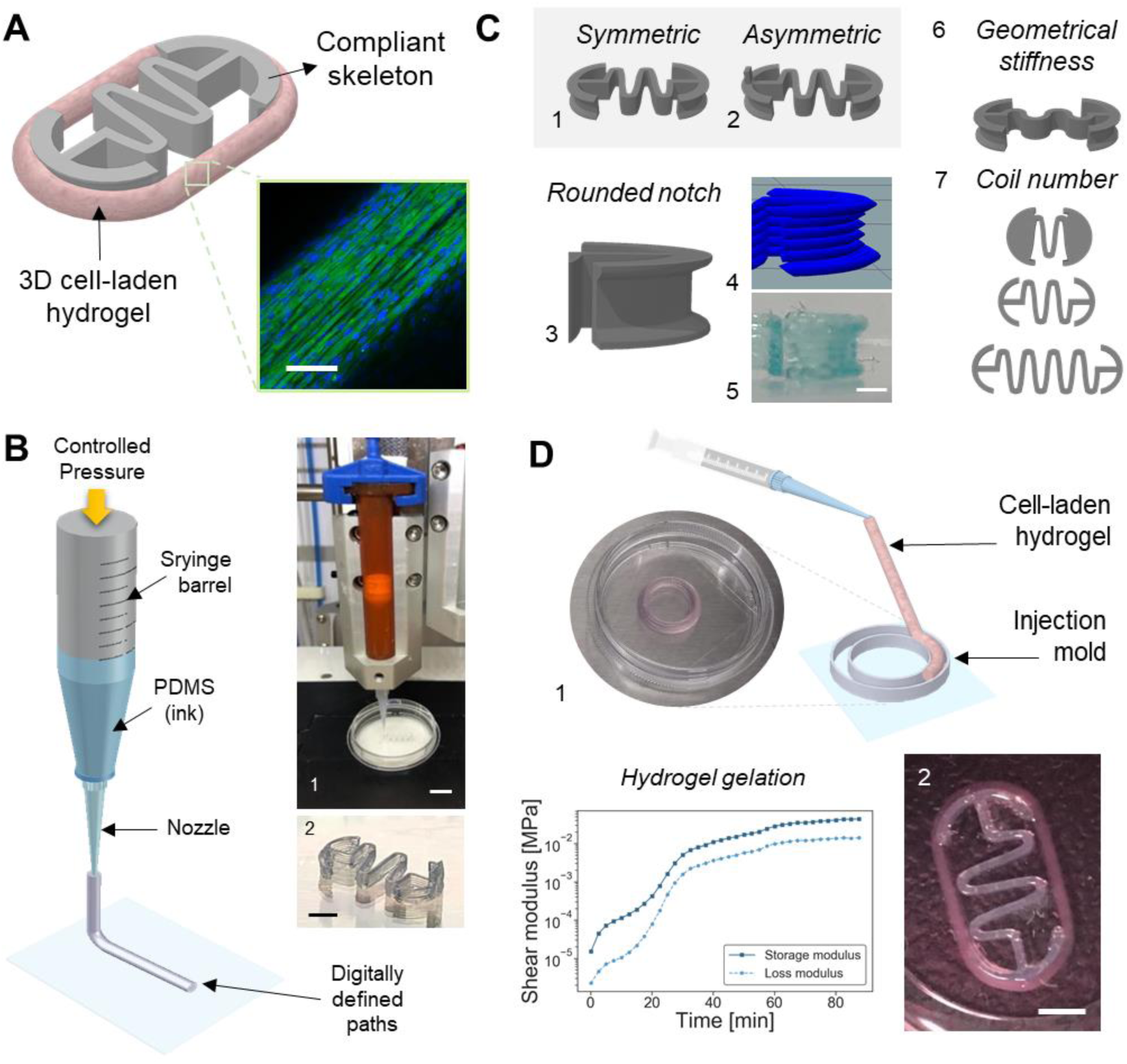
Biobot design and fabrication. (**A**) Schematic of a biobot consisting of a skeletal muscle cell-laden hydrogel acting as a bio-actuator (CLSM image of aligned skeletal muscle cells) assembled to a compliant spring-like PDMS skeleton. Scale, 100 µm. (**B**) 3D printing process based on direct ink writing, where the hydrogel (PDMS) is printed onto a flat surface (1) to obtain the biobot compliant skeleton (image in (2)). Scale bar, 1 cm in (1) and 3 mm in (2). (**C**) Different designs of the compliant skeleton, based on a (1) symmetric or (2) asymmetric serpentine flexure with a (3) rounded notch for a perfect assembly of the cell-laden hydrogel (the (4) 3D visualization of layer-by-layer view of the G-code instructions and (5) image of a printed notch). The effect of (6) different angles on the geometrical stiffness and (7) coil numbers were explored to modulate the mechanical properties of the compliant skeleton design. Scale, 1 mm (**D**) The fabrication process of 3D cell-laden molds (image of a mold (1)) and characterization of (2) mechanical properties related to the of the cell-laden hydrogel cross-linking, as well as a real image of a self-assembled biobot. Scale bar, 3 mm.

Regarding the active biological muscle-based actuator, the cell-laden scaffold was prepared by using customized 3D-printed molds with the desired circular shape and size. A hydrogel composed of fibrinogen, thrombin and Matrigel® and laded with skeletal muscle myoblasts was casted inside the mold (**Fig. 1D**), leaving the whole setup first in growth medium (GM) to let the cells grow and expand, and later on in differentiation medium (DM) to allow the differentiation process ^[49]^. Both GM and DM were supplemented with 6-aminocaproic acid (ACA) to reduce the degradation of the hydrogel due to proteases ^[52]^. The cross-linking of the hydrogel was studied over time to closely evaluate the gelation process and obtain reproducible cell-laden scaffolds for the biobots construction. **Fig. 1D** shows the shear storage and loss moduli of the hydrogel at 37 °C at a frequency of 1 Hz for 90 min. At the start of the characterization, when fibrinogen, thrombin and Matrigel® have been mixed, an initial peak in both moduli after 5 min points at the fast cross-linking of fibrinogen into fibrin by the action of the enzyme thrombin. After approximately 30 min at physiological temperatures, another increase in the absolute value of the shear modulus indicates the thermal-induced cross-linking of Matrigel®. After this point, the structure of the cell-laden hydrogel is stable and warm GM can be added to the injection mold for cell culture.

The preparation and maturation process of the skeletal muscle biobot comprises several stages. **Fig. 2** depicts the process timeline to obtain a myoblast-laden hydrogel. In stage 1, C2C12 cells are embedded in the 3D scaffold and left to grow for three days, leading to a myoblast-laden hydrogel. The cell-laden scaffold is manually transferred to the 3D-printed compliant skeleton based on PDMS in presence of DM (Stage 2). At this point, insulin-like growth factor (IGF-1) present in solution promotes the fusion and differentiation of myoblast into myotubes ^[42]^, supporting the differentiation process and leading to the natural compaction of the muscle-actuator around the compliant skeleton. During the maturation process, the cell-laden scaffold will perfectly adapt to the skeleton’s shape thanks to the rounded edges, avoiding the formation of stresses that could damage the tissue. Due to these compaction forces and to the rounded notches that prevent disassembly, the biobot adopts a buckling structure that provides the necessary asymmetry to the biobots conformation for optimal motion (Stage 3).

**Fig. 2.**
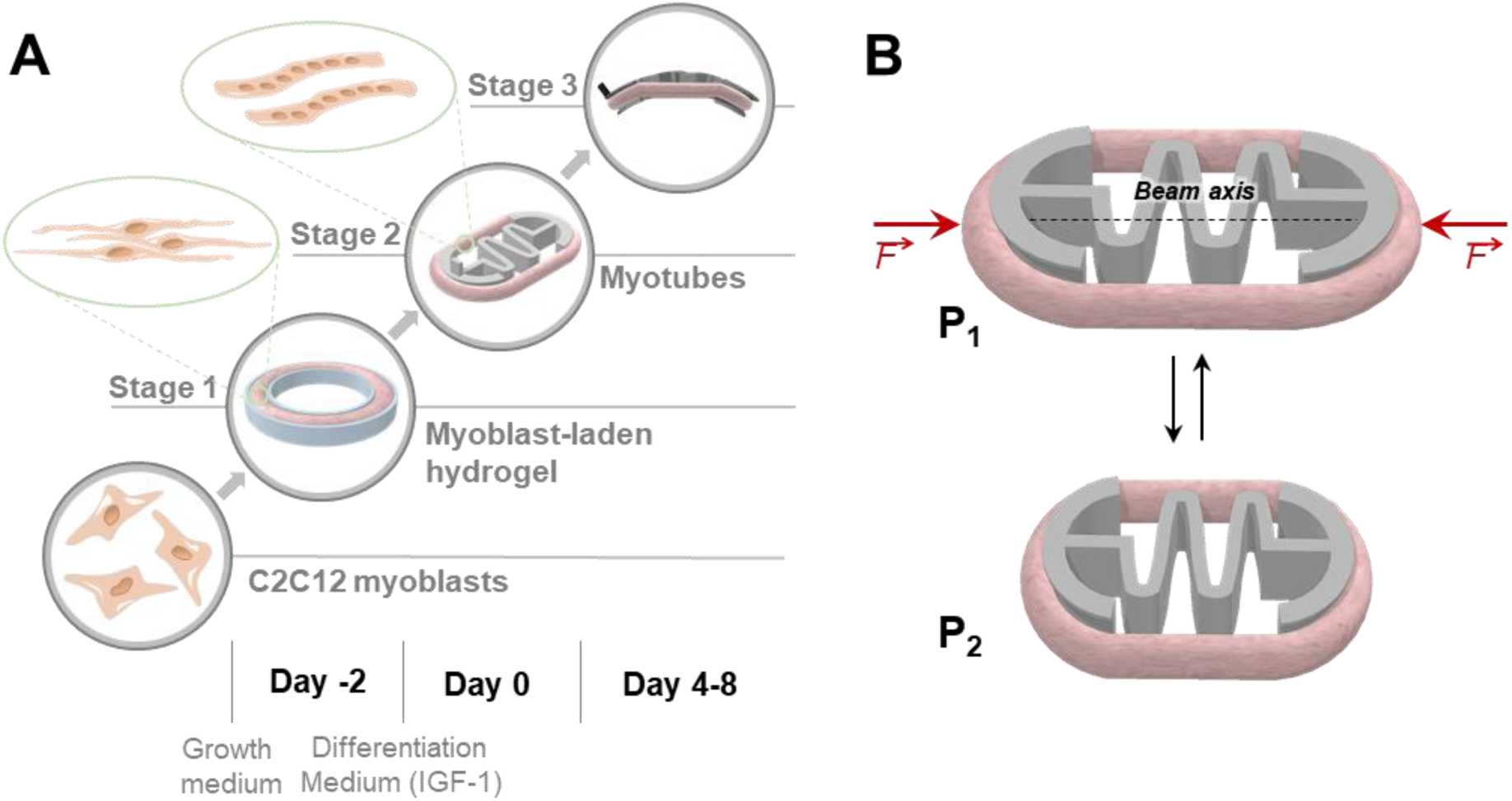
Cell differentiation and compliant mechanism at the biobot platform. (**A**) Conceptual schematic of the cell maturation process at the biobot platform, along with the timeline in which cells seeding, mold preparation (Stage1), assembly (Stage 2), and bending (Stage 3) takes place. (**B**) Schematic representation of the compliant mechanism of the biobot, where compression (P2) and expansion state (P1) of the serpentine flexure are induced by the contraction forces of the muscle and spring-like relaxation behaviour of the assembled muscle cell-based bio-actuator upon EPS.

The compliant mechanism of our untethered bio-hybrid robot is based on the longitudinal force exerted along the beam axis by the muscle tissue around the serpentine-like skeleton upon electrical pulse stimulation (EPS) (**Fig. 2B**). The compliant nature of the spring serpentine structure allows its deformation with low geometrical stiffness and a restoring force that brings the biobot back to its original state.

The skeleton optimal geometrical parameters were evaluated by using finite element analysis (FEA). To study the effect of the spring design on the compression efficiency, we designed three different geometries: i) case 1, with a low-degree variation along the serpentine, creating low amplitude oscillations; ii) case 2, with much larger amplitude in the oscillations, allowing a better distribution of the stresses; and iii) case 3, with large amplitude, but a strict 180° angle at the points of curvature (**Fig. 3A**). In order to mimic the compression from the skeletal muscle tissue contraction on the compliant skeleton, two uniaxial point forces of the same magnitude were applied at both sides of the structure in a 3D simulation (assuming static conditions and only mechanical and linear deformations). The profile of a real twitch contraction was measured by image difference and were normalized in a way that the maximum value was 100 µN (more details in SI section) for a more meaningful simulation of the contraction kinetics. On the right side of **Fig. 3A**, we calculated the von Mises stresses of each skeleton case for a compression force of 100 µN. The von Mises stress is used to predict yielding of materials under complex loading from the results obtained from uniaxial tensile test and provide information about the equivalent stress distribution across the whole structure. The higher and localized stress values obtained for cases 2 and 3 already indicate that these structures can probably compress easier than case 1, although a better characterization of this can be obtained by calculating the geometrical stiffness of the designs.

**Fig. 3.**
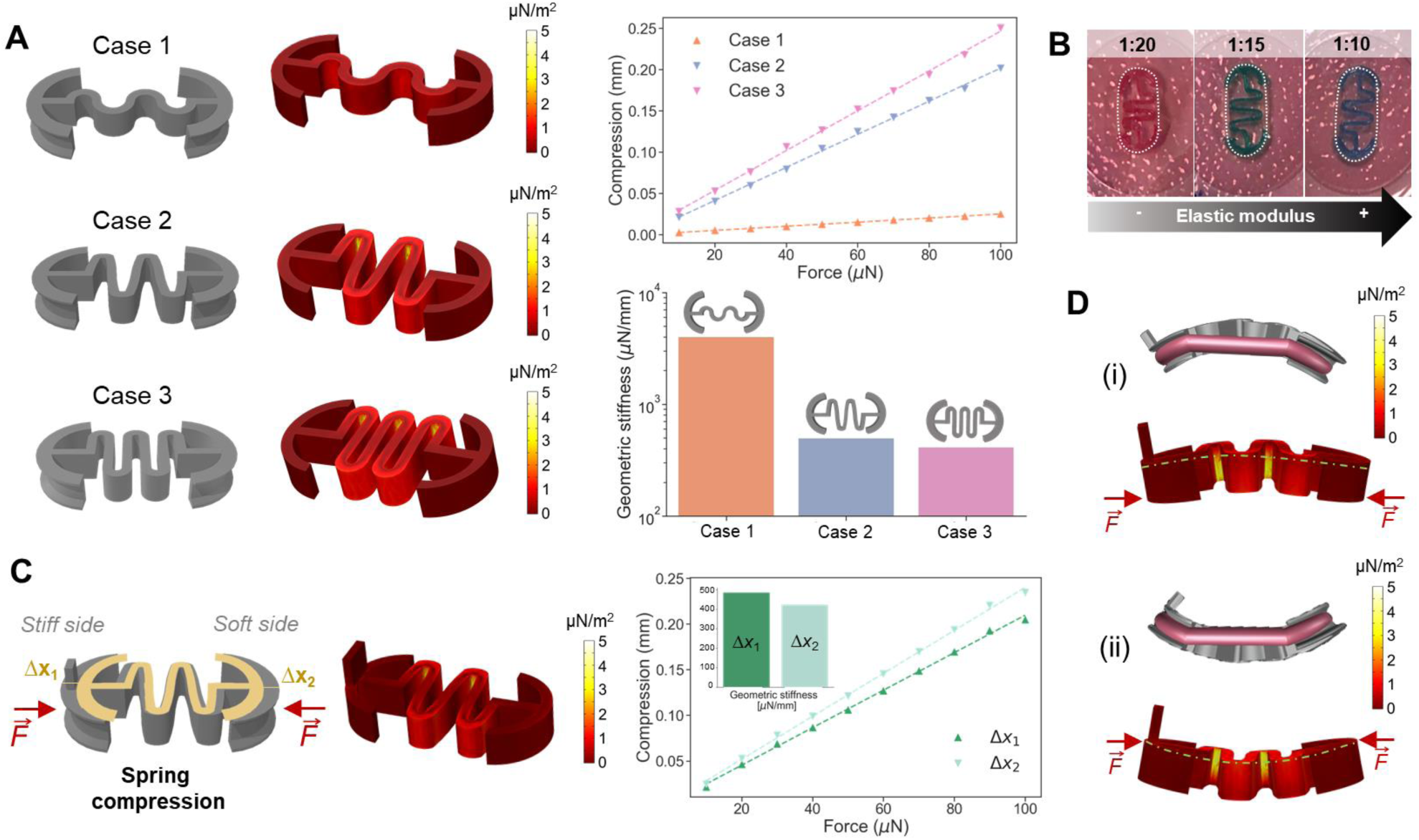
FEA simulations of mechanical deformation on biobot compliant skeleton. (**A**) Three different cases were considered to optimize the coil curvature and shape, depending on the geometrical stiffness of the construct. Next to the 3D representation of each case, the Mises stresses of each structure after a symmetrical load of 100 µN on each side are presented. The maximum compression of the skeleton in terms of the applied force (right top) and geometrical stiffness (right bottom) were obtained per each of the three cases (**B**) Stiffness modulation properties tuned by PDMS chemical composition; different coloured dyes were included to the PDMS to differentiate them, corresponding red, green and blue to 1:20, 1:15 and 1:10, respectively. (**C**) Stiffness evaluation in presence on an asymmetric skeleton. (**D**) Bending analysis allow to study the buckling behaviour on the compliant skeleton, revealing that there is inhomogeneity in the compaction that can be related to buckling, which it can happen both towards the leg or the opposite direction.

The geometric stiffness of the skeletons is a parameter that gives information about the stiffness of the material after applying a deformation, taking only into consideration the geometry of the structure. For each design, we performed a force sweep using the contraction profile with a maximum force in the range of 10-100 µN (symmetrically at both edges), based on reported forces in the literature ^[39,42]^. The maximum uniaxial compression of the compliant skeleton, which coincided in time with the maximum force of the contraction profile, was plotted in terms of the force (left top side **Fig. 3A**), revealing a linear elastic response dictated by Hooke’s law, that allowed us to calculate the geometrical stiffness of the material, *k*, by the equation *F = kx*, where *F* is the applied force and *x* the compression. The inverse of the slope yields the geometrical stiffnesses (left bottom side **Fig. 3A**), where we can see that case 1, with a soft curvature in its coil, had much larger geometrical stiffness than the other two cases, which presented slightly similar values. Further experiments with 3D-printed skeletons revealed that case 3 was easily compressible and collapsed, making the serpentine coils touch and stick to each other. Therefore, the design of case 2 provided the appropriate stiffness conditions for muscle-induced compression in the expected force ranges. However, the compliant structure stiffness can also be tuned by changing the ratio of elastomer to curing agent (**Fig. 3B**). We explored the impact of the PDMS chemical composition by preparing skeletons with the case 2 at different ratio of elastomer to curing agent: 1:20 (red), 1:15 (green) and 1:10 (blue), ordered from less to more stiffness. As expected, the compaction of the cell-laden hydrogel around a skeleton at 1:20 is higher than the 1:10, demonstrating the feasibility to tune the structural configuration of the biobot’ skeleton by changing its mechanical stiffness.

Having defined optimal serpentine structure, the asymmetry of the bio-hybrid robot was also studied by FEA simulations (**Fig. 3C**). Likewise, by simulating two uniaxial equivalent contraction profiles at both sides of the skeleton, we observed that the presence of a small post bulging out of one of the sides induced a differential compression of the structure. In this compression *vs* force plot, it can be seen how the maximum compression of the stiffer side (with the induced asymmetry), termed Δx_1_, was larger than the compression in the softer side, Δx_2_, as demonstrated on the geometrical stiffness evaluation. Finally, we studied the feasibility of the buckling behaviour by inducing a uniaxial compression force at both sides of the skeleton, but out of center, as indicated by the force vectors in **Fig. 3D**. These forces mimicked the passive compaction of the tissue that occurs during myogenesis ^[39,42]^. We hypothesize that this effect could be caused by a spontaneous symmetry breaking during tissue compaction, probably due to a combination of the heterogeneity of the muscle constructs that could lead to an asymmetric distribution of the compression forces and the interaction with close interfaces. For robotics systems at small scales, symmetry breaking is key to achieve efficient motion ^[58]^. Therefore, we expect that this differential compression produces a difference in the fluid flow fields, leading to directional motion.

It is known that dynamic mechanical stimulation is beneficial for the differentiation and maturation of skeletal muscle cells, as it mimics the conditions of native tissue ^[59,60]^. Therefore, we hypothesized that the spring-like configuration of the serpentine skeleton provided dynamic stimulation after spontaneous contractions by reacting with an opposite restoring force that could further expand the tissue, offering mechanical stretching in the form of a feedback loop. To demonstrate this, we compared two types of biobots: i) the compliant and untethered spring-like skeleton, and ii) a 2-post system that was tethered and less compliant (**Fig. 4A**). During myogenesis, we checked for the presence of spontaneous contractions *via* optical microscopy revealing that, after 4 days of differentiation (D4), only spring-like biobots were showing strong spontaneous contractions, fully synchronized and at a frequency of approximately 3 Hz, while bio-actuators in the 2-post system were not showing any spontaneous contractions (**Fig. 4B**). At D8 of differentiation, the muscle tissue in the 2-post system showed small localized contractions that were not synchronized and, eventually, at D10, the spontaneous contractions were strong and globally distributed, as in the bio-hybrid robot (**Fig. 4B, Movie S1**).

**Fig. 4.**
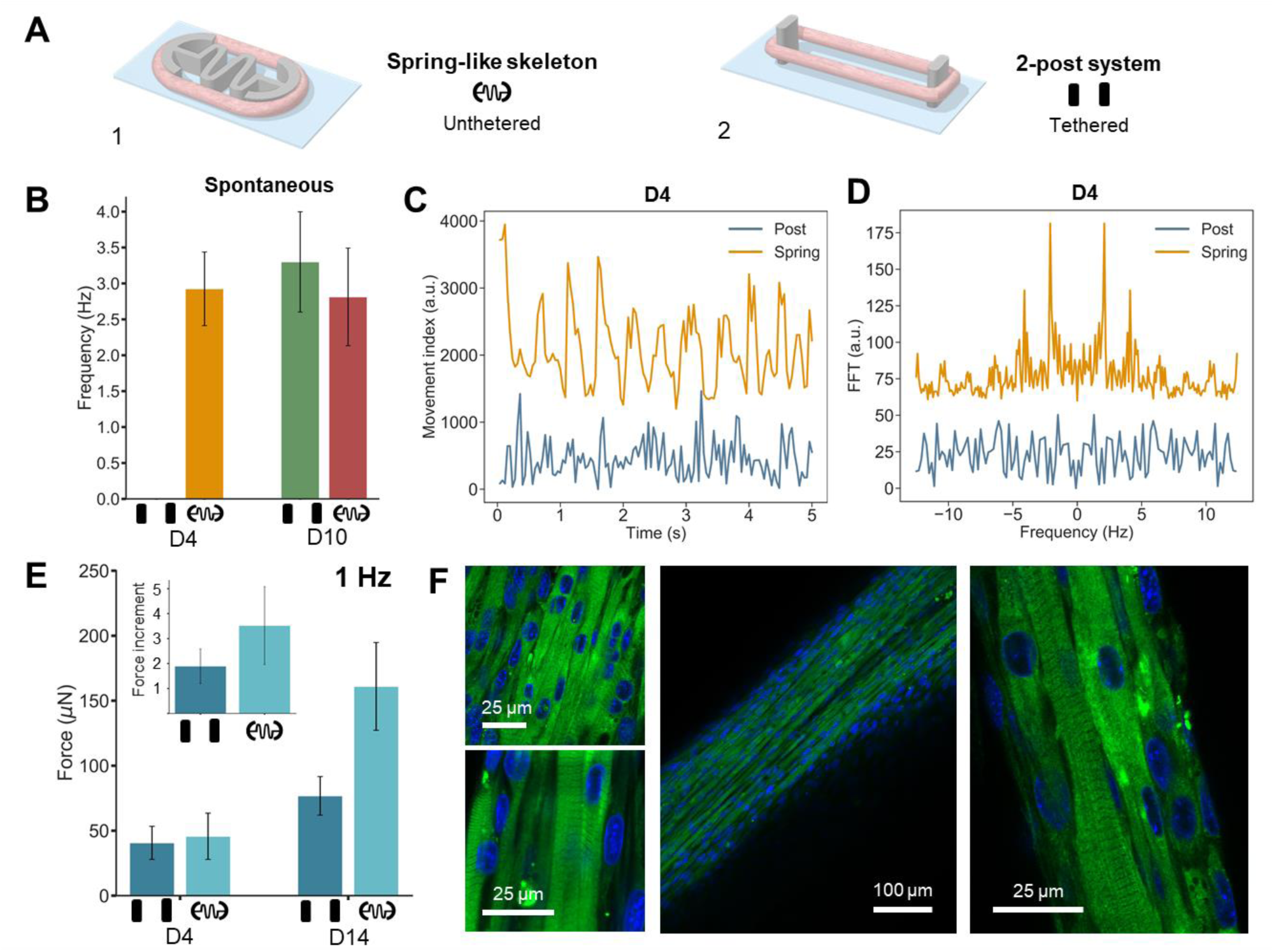
Force generation and biological characterization of skeletal muscle biobots. (**A**) Schematic of spring-like skeleton setup (1) and 2-post system (2). (**B**) Spontaneous contraction evaluation at D4 and D10. N = 4-6 (error bars represent standard error of the mean). (**C**) Movement index and corresponding Fourier transform (**D**) to evaluate spontaneous contraction in 2-post system and spring at D10. (**E**) Force measurement of the 2-post system and biobot system after 4 and 14 days of differentiation, showing an almost four-fold increase in the force (inset) in the latter case, compared to a two-fold increase in the former. N = 3-4 (error bars represent standard error of the mean). (**F**) Confocal evaluation of the tissue ring in a biobot at different magnifications, showing the presence of sarcomeric structures. Myosin Heavy Chain II: green; cell nuclei: blue.

The movement index of the contractions, defined as the pixel difference between images in a small region of interest (ROI) of the tissue, is shown in **Fig. 4D** at D4 of differentiation. It can be seen how only the biobot in a spring-like skeleton showed periodic spikes in its movement index (representing the spontaneous contractions), while the signal of the 2-post system was mainly noise. A fast Fourier transform (FFT) of this signal (**Fig. 4E**) further confirmed that only the former showed synchronized and defined contractions at 3 Hz, while the signal of the latter did not have any defined frequencies and was just noise.

The strength of the contractions was also evaluated and compared (**Fig. 4E, Movie S2**). Although the bio-actuator in the 2-post system did not present spontaneous contractions at D4, EPS could induce contractions in the tissue, which were of a similar magnitude to those induced in the biobot. However, force measurements after several days in differentiation demonstrated that the muscle tissue of the spring-like biobot had increased its force almost 4-fold, while the 2-post bio-actuator only by 2-fold, indicating an enhanced maturation of the tissue (**Fig. 4E**). Confocal immunostaining of myosin heavy chain II (MyHCII) and cell nuclei showed well aligned myotubes with the presence of sarcomeric structures in the hybrid biobot (**Fig. 4F**).

After at least 4 days of differentiation, the hybrid biobots could show a spontaneous symmetry breaking leading to a buckling structure, as depicted in **Fig. 2A** and validated in **Fig. 3D**. However, due to the hydrophobicity of PDMS and surface tension at the air-liquid interface, biobots with a symmetric skeleton remained floating without showing any buckling behaviour (**Fig. 5A1, Movie S3**). An asymmetry in the form of buckling was induced in the symmetrical skeletons by forcing the biobot inside the culture medium, while working in a plastic Petri dish, during the differentiation process.

**Fig. 5.**
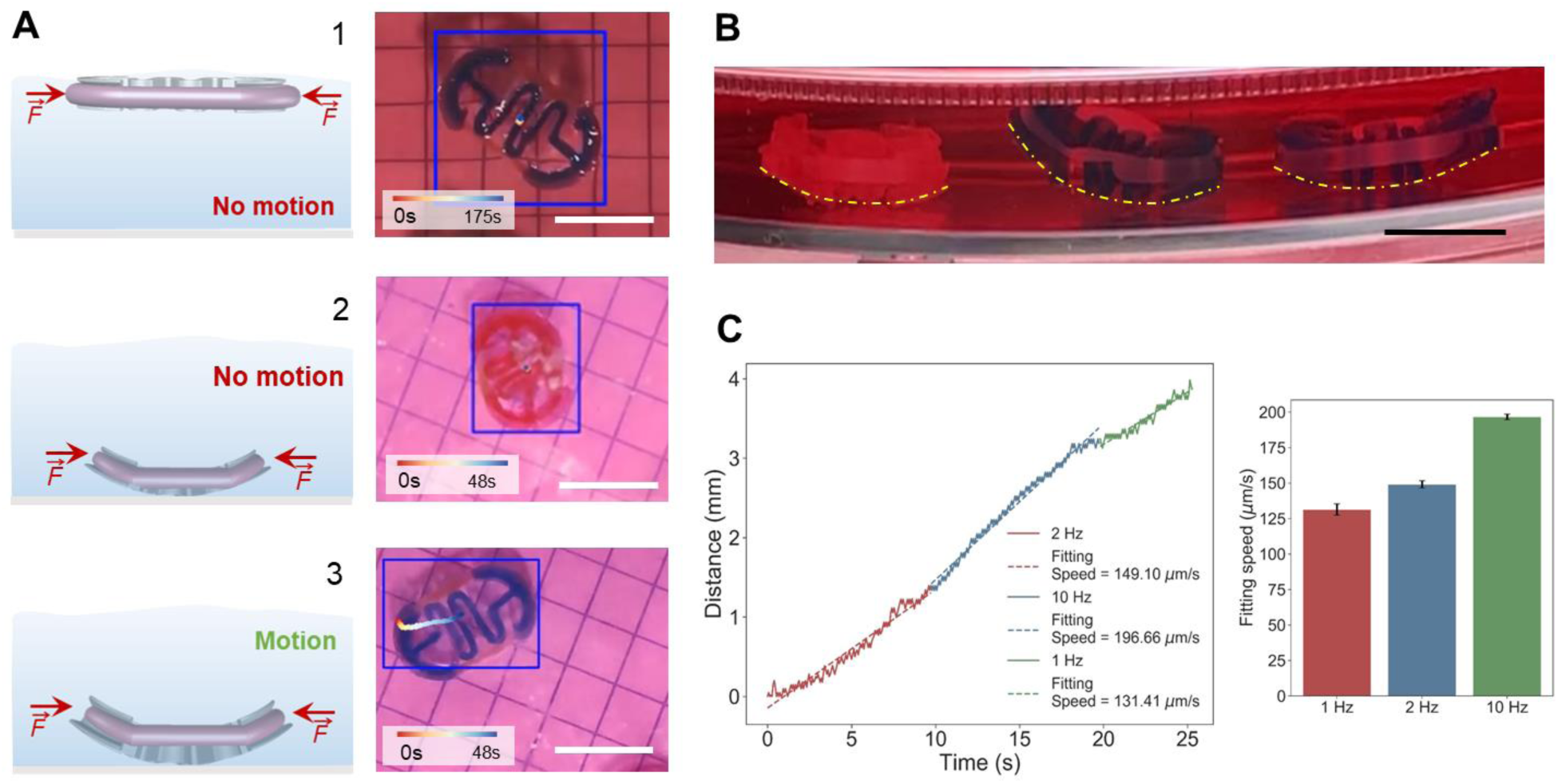
Symmetric biobot motion evaluation. (**A**) Schematic of the side position and top image from the symmetric biobot when located at the interface (1), and in solution for the two cases where no motion (2) and motion (3) is taking place. Scale bar, 10 mm. (**B**) Side image of three different biobots with different stiffnesses, with the corresponding bending due to the buckling effect. Scale bar, 10 mm. (**C**) Speed analysis for the case of the symmetric biobot at different frequencies. Error bars represent the error of the least-squares fitting.

For the biobots with a symmetric skeleton, two different cases where studied. In the first one, considered the biobot, maintained at the air-liquid interface, no significant motion was observed (**Fig. 5A1, Movie S3**). However, when the symmetric biobot was placed near to the bottom surface, buckling deformation was spontaneously generated due to the heterogeneity of the muscle construct yielding a net compaction force outside its axis of symmetry (**Fig. 5A2 and 5A3**), observing motion for some of the symmetric biobot upon EPS. This spontaneous symmetry breaking and thus the degree of deformation could not be controlled, but it was dependent on the stiffness of the material, given by the curing-agent-to-base ratio of PDMS. In **Fig. 5B**, we can see that low stiffness PDMS (left) would completely compress and collapse the structure, while higher stiffnesses (medium and right) would allow some freedom of movement that could result in net motion. **Fig. 5C** shows an example of the displacement of the best-case scenario of symmetric swimming, which could reach maximum speeds of 100-200 µm/s (or 0.5-1 bl/min), increasing with frequency (**Movie S4**). This type of motion could resemble the swimming style of certain fish near surfaces, such as the burst-and-coast behaviour of zebrafishes, characterized by sporadic bursts followed by coasting phases ^[61,62]^. The strong hydrodynamic couplings between the flow field around the biobots and the bottom surface, which are known to play a substantial role in the motion of microorganisms ^[63]^, may also play a significant part in this case. For instance, it is known that hydrodynamic coupling induces the alignment of the swimming direction of a microorganism with the nearby surface ^[63–66]^. It should be considered, however, that the motion resulting from symmetric skeletons was not predictable, as it strongly depended on the degree of buckling curvature, which could not be controlled. Moreover, both the speed and the direction of motion were not clearly defined, as they relied on spontaneous symmetry breaking of the structure and its interactions with the surfaces. Therefore, consistent and controllable motion was only obtained when an asymmetry was previously incorporated in the design.

Motion with asymmetric skeletons proved to be predictable in terms of yield and direction of swimming. The presence of the post on one of the sides of the skeleton’s design induced a different stiffness of the two sides of the skeletons, as previously demonstrated by FEA analysis in **Fig. 3C**, as well as allowing stable floating of the buckled structure, unlike in the case of the symmetric biobots that required to be placed to the bottom surface (**Fig. 6A**). In general, while symmetric designs moved only under certain conditions that broke its symmetry at speeds lower than 100 µm/s, asymmetric biobots swam at much higher speeds and in a consistent manner (direction opposite to the post), although with great variability between samples (**Fig. 6B**).

**Fig. 6.**
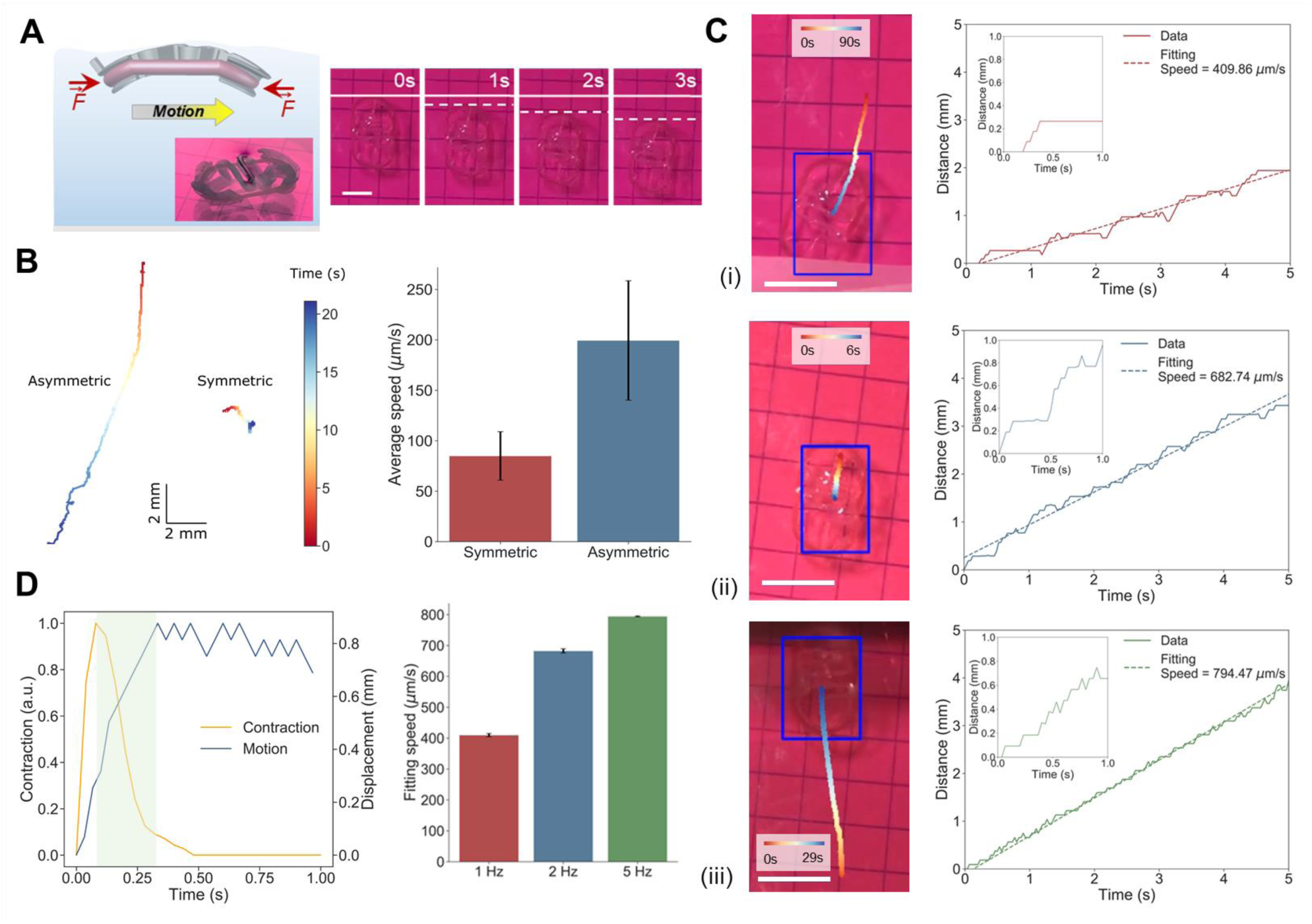
Asymmetric biobot evaluation. (**A**) Schematic of the side position and time step image of an asymmetric biobot motion for 3 seconds and its corresponding speed under different EPS frequencies. (**B**) Average speed of symmetric and asymmetric biobots, and the track motion comparison of an individual symmetric and asymmetric biobots. N = 8 (symmetric) and N = 18 (asymmetric). Error bars represent standard error of the mean. (**C**) Motion evaluation of the asymmetric biobot, depicting its motion efficiency with an image where the corresponding track is shown, as well as its displacement over time, at (i) 1 Hz, (ii) 2 Hz, and (iii) 5 Hz. Scale bar, 10 mm. (**D**) Superposition of a measured contraction (yellow) with the motion of a biobot after one contraction. The green-shadowed portion represents the region of motion that can only be explained by inertia. Average motion of asymmetric biobots upon different EPS frequencies. Error bars represent the error of the least-squares fitting.

Three tracking examples for the biobots with an asymmetric skeleton at different stimulation frequencies are also shown, where speeds of more than 700-800 µm/s for frequencies of 2 Hz and 10 Hz, and 550 µm/s for 1 Hz can be observed (**Fig. 6C, Movie S5**). In the insets of 1 Hz and 2 Hz, we can see how the motion occurs in a stepwise manner, while for 5 Hz it is more continuous. Comparing the average trajectories of both designs from all the trackings, we found that both symmetric and asymmetric biobots were able to achieve motion. An interesting common feature of asymmetric biobots is the stepwise motion, consistent with swimming mechanism driven by inertia (**Fig. 6D**). In fluid mechanics, the relationship between inertial forces and viscous forces during the motion of a swimmer is known as the Reynolds number (Re). This dimensionless quantity allows to differentiate between regimes of motion in which laminar flows (typical of viscous motion) or turbulent flows (typical of inertial motion) are dominant. At low Re (≪ 1), where viscous forces dominate, the fluid dynamics are described by the time-independent Stokes equation, in which inertial components are considered negligible. At this scale, the “scallop theorem” by Purcell dictates that a swimmer must perform non-reciprocal or time-irreversible motion to achieve a net displacement different from zero ^[67]^. Microorganisms manage to break this time-reversal symmetry by rotatory motions ^[68]^, like those of bacterial flagella ^[69]^, which have been mimicked by artificial micropropellers ^[70]^ or bio-hybrid swimmers based on cardiac cells ^[71]^. In our case, the compression mechanism of the skeletal muscle tissue against the skeleton is time-reversible since the shape changes are identical if time is reversed. This bio-swimmer, therefore, should not present motion at low Re. The Re number is defined as *Re* = *v L*/*ν*, where *v* is the characteristic fluid velocity, *L* is the characteristic length of the swimmer and *ν* the kinematic viscosity of the fluid. In our case, given an approximated size of 10 cm and speeds of the biobots between 100-500 µm/s, we find Re numbers of the order of 1-5. In this range, both viscosity and inertia play a significant role, and the motion cannot be considered neither purely viscous nor purely inertial.

Hydrodynamics simulations were performed to demonstrate that the fluid inertia and the asymmetric deformation of the biobot upon muscle contraction are key to its locomotion. Due to the high computational power of simulating the deformation of three-dimensional structures coupled with hydrodynamics, a 2D model was used instead (see SI). As shown schematically in **Fig. 7A**, we modelled the time-dependent deformation of the left side of the biobot as *d*_*l*_(*t*)= Δ_*l*_ *g*(*t*), and of its right side as *d*_*r*_(*t*)= Δ_*r*_ *g*(*t*). The deformation of the skeleton in the middle was set to vary linearly between these two values (see SI). The maximum compressions, Δ_*l*_ and Δ_*r*_, were equal in the case of a symmetric biobot but different in the case of an asymmetric one with distinct geometric stiffness on each side. The periodic deformation of the skeleton due to the contraction and the relaxation of the muscle cells, *g*(*t*), was chosen to closely follow the measured deformation in the experiments, as shown in **Fig. 7B**. This contraction profile is typical of skeletal muscle tissue, as it is known to go through a fast period of contraction followed by a slower relaxation (**Fig. 6D**) ^[39]^. Since the contraction and relaxation of the skeleton are time-reversible we expected propulsion only if Δ_*l*_ is different from Δ_*r*_.

**Fig. 7.**
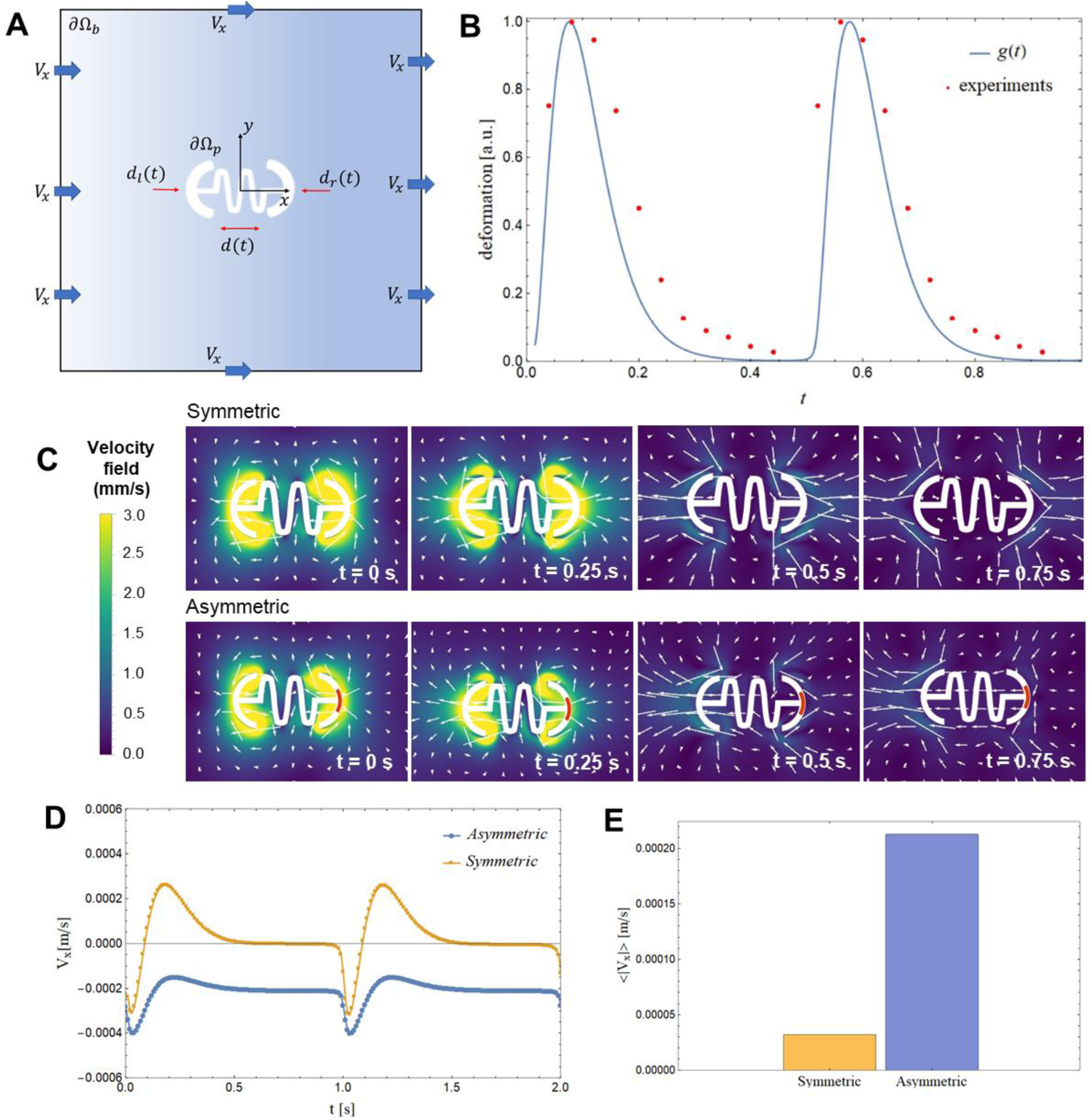
Hydrodynamics of biobots locomotion. (**A**) Schematics of the 2D computational model. (**B**) Periodic deformation of the biobot used in the simulation compared to that measured in the experiments. (**C**) Magnitude of the velocity field and streamlines at four different instants during one period, in the case of a biobot driven at 1Hz. The top panels show the result for a symmetric biobot, the bottom panels show the results for an asymmetric biobot. The orange icon at the right side of the biobot represents the region with the post (stiffer section experiencing smaller deformations). (**D**) Evolution of the x component of the velocity of the biobot during two periods. (**E**) Average speed of the biobot over one period in the case of a stimulation at 1Hz.

The snapshots in **Fig. 7C** show the magnitude of the velocity field and the streamlines around the biobot during one contraction/relaxation period as computed from the numerical simulations. In the top panels, displaying the symmetric case, the left and the right side of the biobot deformed in the same way Δ_*r*_= Δ_*l*_. As a result, the streamlines on the left and on the right of the biobot were highly symmetric and the net displacement of the biobot, averaged over one period, was negligible. This is confirmed by looking at **Fig. 7D** and **E**, which show the velocity of the symmetric biobot as a function of time and its average over one period.

In contrast, a biobot with a stiffer right side deformed asymmetrically, undergoing a larger deformation along its left side Δ_*l*_>Δ_*r*_. In the bottom panels of **Fig. 7C** we show snapshots of the magnitude of the velocity and of the streamlines during one period for an asymmetric biobot. The streamlines at t = 0.5 s and t = 0.75 s indicate that the biobot is moving from the right to the left (right side of the skeleton corresponding to the stiffer side), which is in agreement with what observed in the experiments. This finding is confirmed in **Fig. 7D** where the velocity of an asymmetric biobot is shown to be negative, i.e. from the right to the left, during the entire period. The average speed of the asymmetric biobot over one period is compared to the symmetric one in **Fig. 7E**, confirming our hypothesis that an asymmetric deformation of the biobot is necessary for its locomotion.

## Discussion

Biological robots based on skeletal muscle tissue are of interest in the soft robotics area due to the inherent properties of natural tissue that are difficult to replicate with artificial actuators. Muscle-based biobots have been proposed not only as tissue models to study muscle development and regeneration ^[39,72]^, or as drug testing platforms in the biomedical field ^[73,74]^, but these actuators are posed as excellent candidates to study the integration of tissue with complex artificial materials to understand or improve motion mechanisms through biomimetic approaches in robotic platforms ^[37,38,41,42]^. Here, we report a bio-hybrid robot based on skeletal muscle cells that can swim at speeds comparable to their cardiac counterparts (**Table 1**), also taking advantage of the adaptability and control capabilities inherent to skeletal muscle cells. We reported a biological robot based on a flexible serpentine spring design that is optimized through simulations and subsequently 3D printed. The novelty of a compliant spring-like scaffold instead of a stiffer or tethered one lies in the improved differentiation of the tissue through mechanical self-stimulation upon spontaneous contractions, which creates a feedback loop due to the restoring force of the spring. Moreover, the asymmetry from the serpentine spring skeleton allows the structure to move with two different modes: by inertial swimming at the interface or coasting near a bottom surface. Serpentine spring structures are well-known in the microelectromechanical systems (MEMS) field to create flexible structure by working with microfabrication techniques ^[75,76]^. Although serpentine springs have been implemented in untethered micro-force sensing microrobots to obtain compliant structures ^[77]^, such structures have not been included before in a soft robotic living system. The design of the serpentine spring was ideal to be printed by additive manufacturing, obtaining a 3D structure of attractive properties from the mechanical point of view, being of great interest for the design of future 3D printed robotic platforms.

The optimization of the designs, based on several features like asymmetry or coil curvature, was performed *via* FEA simulations, which allowed us to find the appropriate geometrical stiffnesses of each case and then experimentally test them. As expected, we found that the addition of an asymmetric feature in one of the sides resulted in a differential compression of the skeleton, which is known to be necessary for efficient motion ^[42]^. Moreover, force measurements revealed that differentiation in a compliant spring-like skeleton was beneficial when compared to other static tethered skeletons, such as a 2-post system. The muscle tissue under these conditions showed earlier spontaneous contractions and greater increments of force during the differentiation process. We hypothesize that the dynamic compliance of the skeleton, reacting to the compression with a restoring force, provided an additional level of cyclic mechanical stimulation that helped to achieve a better degree of differentiation. In fact, the potential of expanding the use of such skeleton in other muscle-based biobots configurations would represent a novel approach to implement tailored training protocols that do not require for any external stimuli.

Finally, we characterized the motion of two types of bio-hybrid robots: a symmetric swimmer and an asymmetric swimmer. For the former, we found that motion at the air-liquid interface was not possible due to the generated symmetric flow fields, which was supported by hydrodynamic simulations. In contrast, when the symmetric biobots were placed into the culture medium, a buckling behaviour was observed and the swimmer adopted a curved structure that allowed motion at low speeds, although such motion was unreliable, most likely due to interactions between the generated flow fields and the surface. Asymmetric biobots, however, presented a reliable and consistent motion, as they already displayed buckling when floating in the air-liquid interface. Stimulation with EPS showed high speeds, up to 800 μm/s and motion studies, as well as hydrodynamic simulations, were consistently confirming the hypothesis that inertia plays a key role in the motion mechanism. Future work should aim at understanding the interactions between the skeleton and surfaces, to comprehend the exact parameters governing this type of motion and cooperative behaviour of several biobots.. Moreover, the scalability of these biobots could be investigated by comparing the motion of swimmers with different number of coils and exploring other configuration, mainly due to the versatility that 3D bioprinting offers in terms of shape and multi-material 3D printing, as well as studying the implementation of nanocomposities.

In summary, the bio-hybrid swimmer based on skeletal muscle cells hereby reported move at speeds faster than the largest skeletal-muscle-based biobots up to date ^[54]^ and comparable to other cardiomyocyte-based bio-swimmers ^[37]^. We use 3D printing of PDMS to fabricate serpentine-like skeletons that can act as a spring when a muscle ring compresses them, resulting in compliant scaffolds that aid in the differentiation of the tissue. The optimization of the designs via FEA simulations allowed us to find the appropriate geometrical stiffnesses by studying their asymmetry and coil curvature. We found that differentiation in a compliant spring-like skeleton was beneficial when compared to other static tethered skeletons due to its associated mechanical self-training capabilities, resulting in earlier spontaneous contractions and greater increments of force during the differentiation process. We hypothesize that the dynamic compliance of the skeleton, reacting to the compression with a restoring force, provides an additional level of cyclic mechanical stimulation that aids to achieve an improved maturation. Also, the hydrodynamics study of the different motion modes provides a fundamental understanding over its motion mechanism, demonstrating the potential of applying such serpentine-like structure to provide asymmetry and achieve enhanced force outputs on future biobots configurations. Further research in the bio-hybrid robotics field should be focused on the integration of several tissues and obtaining more complex, yet useful, ways of actuation that can finally prove the benefits of using native muscle tissue instead of man-made soft actuators. Once achieved, the next main challenges will reside in ensuring long-term stability of the constructs and tolerance to different environments, requiring the implementation of novel and soft bioreactors to protect the tissue and the improvement of the control mechanisms. All these advances will undoubtedly imply a coordinated interdisciplinary effort between different fields of expertise, ranging from 3D bioengineering and biology, to materials science and mechanical engineering.

## Materials and Methods

### Biobot fabrication

C2C12 mouse myoblasts were purchased from ATCC. Growth medium (GM) consisted of high glucose Dullbecco’s Modified Eagle’s Medium (DMEM; Gibco®) supplemented with 10% Fetal Bovine Serum (FBS), 200 nM L-Glutamine and 1% Penicillin/Streptomycin. Cells below passage 4 were used before reaching 80% confluency in Corning® T-75 flasks. Differentiation medium (DM) consisted of high glucose DMEM containing 10% Horse Serum (Gibco®), 200 nM L-Glutamine (Gibco®), 1% Penicillin-Streptomycin (Gibco®), 50 ng/ml insulin-like growth factor 1 (IGF-1, Sigma-Aldrich) and 1 mg/ml 6-aminocaproic acid (ACA, Sigma-Aldrich).

C2C12 skeletal myoblasts were harvested from the flask when reaching 80% confluency. A centrifuged cell pellet with 3 million cells in total was mixed with 54 µL of cold GM (32.5% v/v) supplemented with 1 mg/mL of ACA (Sigma-Aldrich), 12 µL of a stock solution of 50 U/mL of thrombin (7.5% v/v), 90 µL of Matrigel® (60% v/v) and 150 µL of a stock solution of fibrinogen at 8 mg/mL (50% v/v), in that order. The mixture was manually casted immediately on a 3D-printed PDMS injection mold. The tissue construct was left in a cell incubator (37 °C, 5% CO_2_) for 30 min, and GM supplemented with ACA to avoid degradation of collagen by proteases was added and kept for 2 days. Then, the tissue was gently lifted from the mold and transferred to either a (i) 3D-printed compliant skeleton or (ii) a 2-post system. Spring like skeletons of different ratios (1:20, 1:15 or 1:10, with or without dye) were cured at 80 °C overnight. The designs of the skeletons were done with AutoCAD (v. 2019), exported as .stl files, and transformed into GCode, to be later printed with Cellink’s Inkredible+ 3D bioprinter. The 2-post system (3 mm high, 0.5 mm wide and with 2 mm of lateral width) was 3D-printed with PDMS of a 1:20 and crosslinked at 37 °C for several days. Once cell-laden scaffold is transferred to the compliant skeleton, the culture medium was changed to DM supplemented with ACA and IGF-1. After 4 days in DM, the structure is drawn in a plastic petri dish to obtain the desired buckling effect.

### Structural mechanical analysis from the compliant skeleton

Optimal geometrical parameters on the compliant skeleton were evaluated by using finite element analysis (FEA). By performing a force sweep in the range of 10-100 µN (symmetrically at both edges), based on previous reports ^[39,42]^, we can simulate a compression of the skeleton in the same way as the tissue would do. The maximum compression of the complaint skeleton was plotted in terms of the force (**Fig. 3**), revealing a linear elastic response dictated by Hooke’s law, that allowed us to calculate the geometrical stiffness of the material, *k*, by the equation *F = kx*, where *F* is the applied force and *x* the compression. With small forces in the range of hundreds of µN, the yield criterion of the von Mises stress is not fulfilled, as the maximum stress is much lower than the yield stress of PDMS ^[78]^. Therefore, a linear analysis is appropriate to model the deformations of the material.

### Electrical pulse stimulation

The setup for EPS was composed by a waveform generator (PM8572, Tabor Electronics), an oscilloscope (DS1104Z, Rigol), a signal amplifier x15, and a set of carbon-based handmade electrodes consisting of two graphite rods (cat. number 30250, Ladd Research) placed on opposite sides of a plastic Petri dish (**Fig. S2**). Both for force measurement and motion evaluation experiments, the recording was carried out inside an inverted microscope (DMi8, Leica) in a chamber that allowed to mimic physiological conditions (37 °C and 5% CO_2_). Motion videos of the biobots were recorded with a smartphone, keeping a grid paper below the Petri dish for calibration purposes. Pulses at a different frequency and constant width (1 ms) and voltage (15 V) were applied to study its effect on the biobots motion performance.

### Biobot force characterization

The protocol described by Mestre *et al*. ^[39]^ for force measurements in a two post-system for muscle-based actuators was adapted to determine the role of the compliant skeleton on the fabricated biobots’ final force. The recording of the whole setup undergoing EPS was carried out inside an inverted microscope (DMi8, Leica), in a chamber that allowed to mimic physiological conditions (37 °C and 5% CO_2_). For force measurement of the biobots in **Fig. 4**, the tissue was gently transferred at D4 or D14, depending on the experiment, into a 2-post system, and their force measured by deflection of the posts by the cell-laden scaffold. Pulses of different frequencies of 1 ms were applied, keeping a constant voltage of 15 V. Euler– Bernoulli’s beam bending equation was used to estimate the forces and stresses exerted against the posts to the tissue during electrical stimulation ^[39]^ (more details in SI).

### Motion analysis

Motion evaluation of the bio-hybrid robot was performed with a homemade Python tracking script based on computer vision algorithms that could characterize the motion of the biobots after being recorded with any type of smartphone camera (**Fig. 5**). In brief, the video file of the recorded biobots was loaded into the script and the first frame was prompted for the user to manually select an ROI covering the whole biobot area (**Fig. S1**). Then, an object tracking algorithm based on an online AdaBoost feature selection ^[79]^ was applied to this ROI through every frame of the video (more details in SI).

### Immunostaining

Tissue constructs were washed three times in PBS and then fixed with a 3.7% paraformaldehyde in PBS for 15 min at RT, followed by three washes in PBS and stored until use. For immunostaining, cells were permeabilized by 0.2% Triton-X-100 in PBS. After washing thrice in PBS, the constructs were incubated with 5% Bovine Serum Albumin (BSA) in PBS (PBS-BSA) to block unspecific bindings. Then, the tissues were incubated for 2 h at RT and dark conditions with a 1/400 dilution of Alexa Fluor®-488-conjugated Anti-Myosin Heavy Chain II antibody (eBioscience) in 5% PBS-BSA. The unbound antibodies were washed out with PBS, and cell nuclei were stained with 1 µl/mL Hoechst 33342 (Life technologies). Finally, the tissue samples were washed thrice in PBS and stored at 4 °C until their analysis. The fluorescently labelled tissue constructs were imaged under a Zeiss LSM 800 confocal scanning laser microscope (CSLM), with a diode laser at 488 nm and 405 nm excitation wavelength for Myosin Heavy Chain II and cell nuclei.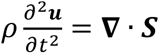

### Statistical analysis

Experiments from spontaneous contractions and force measurement in **Fig. 4** were performed for N = 3-6 independent repeats and error bars represent standard error of the mean. Motion fittings in **Fig. 5** and **Fig. 6** were performed with the least squares fitting method to a linear equation of the form *x = v ·t*, from which the speed was extracted. Average speeds of **Fig. 6B** come from N = 8-18 replicates. When a single biobot was chosen for an in-depth analysis, the error bars in speed bar plots correspond to the error of the fitting.

## Acknowledgments

M.G. thanks MINECO for the Juan de la Cierva fellowship (IJCI2016-30451), the Beatriu de Pinós Programme (2018-BP-00305) and the Ministry of Business and Knowledge of the Government of Catalonia. M.G. and G.Z. thanks Barcelona Institute of Science and Technology for the BIST Ignite Grant (ElectroSensBioBots). R.M. thanks “la Caixa” Foundation through IBEC International PhD Programme “la Caixa” Severo Ochoa fellowships (code LCF/BQ/SO16/52270018). T.P. thanks the European Union’s Horizon 2020 research and innovation program, under the Marie Sklodowska-Curie Individual Fellowship (H2020-MSCA-IF-2018, DNA-bots). M. D. C. acknowledges funding from the European Union’s Horizon 2020 research and innovation program under the Marie Skłodowska-Curie action (GA 712754), the Severo Ochoa programme (SEV-2014-0425), and the CERCA Programme/Generalitat de Catalunya. S.S. acknowledges the CERCA program by the Generalitat de Catalunya, the Secretaria d’Universitats i Recerca del Departament d’Empresa I Coneixement de la Generalitat de Catalunya through the project 2017 SGR 1148 and Ministerio de Ciencia, Innovación y Universidades (MCIU) / Agencia Estatal de Investigación (AEI) / Fondo Europeo de Desarrollo Regional (FEDER, UE) through the project RTI2018-098164-B-I00. This project was also partially funded by Agencia Estatal de Investigación (CEX2018-000789-S).

## Author contributions

M.G, R.M and T. P. conceived the study design, performed the experimental work and corresponding data collecting and analysis, and manuscript writing. G.Z. participated in the experimental procedures. The mechanical and hydrodynamical simulations on finite element analysis were conducted by R. M. and M. D. C., respectively. M. D. C. also contributed to the manuscript writing. S.S. participated in the study design and manuscript writing.

## Data and materials availability

All data needed to evaluate the results and conclusions are included in the main text or the Supplementary Material.

