## Supplementary Information for "Bio-hybrid soft robots with self-stimulating skeletons"


### SUPPLEMENTARY MATERIALS

#### Movies at Supplementary information

**Movie S1:** Spontaneous contraction evaluation between the 2-post system and the spring-like.

**Movie S2:** Force evaluation between the 2-post system and the spring-like biobot.

**Movie S3:** Spring-like symmetric biobot actuated at the air-liquid interface (control).

**Movie S4**: Spring-like symmetric biobot at the bottom surface actuated at different frequencies.

**Movie S5:** Spring-like asymmetric biobot at the air-liquid interface actuated at different frequencies.

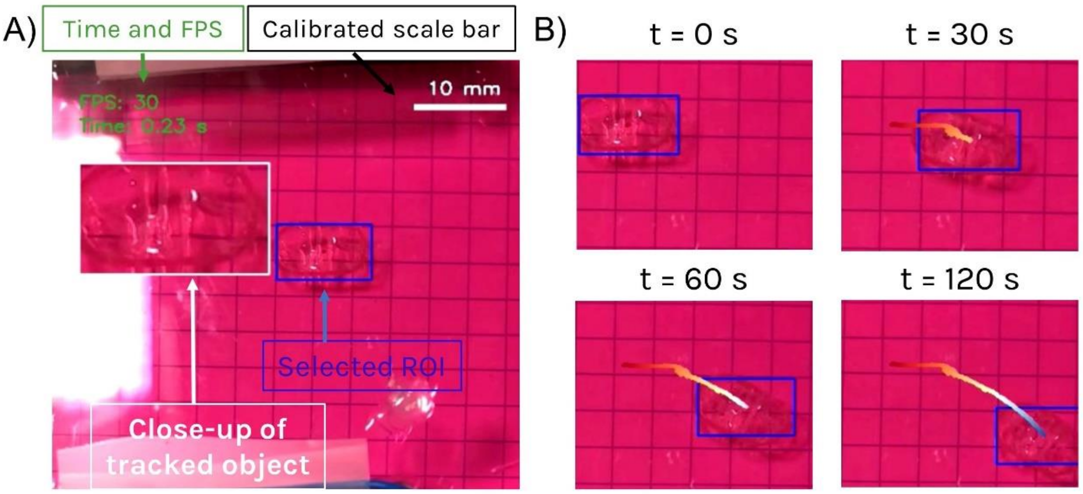


**Fig. S1**: Home-made script for tracking of bio-bots. (**A**) Snapshot of the initial frame of the tracking, in which several features can be observed: i) the time and FPS of the video are stamped on the top left corner of the video; ii) an ROI is manually selected to contain the biobot, or part of it, which is tracked along the video; iii) the grid paper below the Petri dish is used to calibrate the spatial dimensions; and iv) a close-up of the tracked object inside the ROI is displayed to make sure that the biobot is being properly tracked. (**B**) Snapshots of a tracking, showing a color-coded trajectory, going from red (short time) to blue (long time).

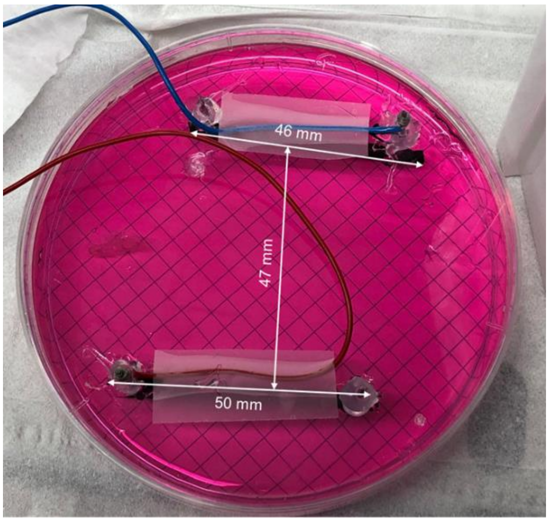


**Fig. S2**: Electrical pulse stimulation (EPS) setup. The experiments were run in a plastic petri dish of 8.8 cm diameter, where the electrodes where coupled after the biobot was moved to the cultures media and conveniently located at the interface or inside the solution.

### Object tracking algorithm

The bio-bot tracking algorithm was written in Python (v. 3.7) and was based on machine learning through an online AdaBoost feature selection algorithm that was applied to a ROI across every frame of the video. This algorithm extracts identifying features from the object to track its position through time, even if the biobot rotates and changes direction. As smartphone videos can be located at different distances and are not calibrated, the script also allowed manual calibration of the video by using a calibrated grid paper as background. Before starting the tracking, the user could manually draw a line along one of the squares and the script would find the conversion pixel/mm to calibrate the displacements and create a scale bar at the top of the image. Moreover, time in seconds and frames per second (FPS) of the original video were displayed on top, as well as a close-up image of the tracked object online, to ensure that the tracking was capturing the bio-bot properly during the video. The central position of the ROI in mm was stored and used to generate a color-coded trajectory on the images (**Fig. S1**), as well as to plot the total displacement and compute the speed of motion.

**Biobots Reynolds number regime**

The Reynolds number (Re) allows to differentiate between regimes of motion in which laminar flows (typical of viscous motion) or turbulent flows (typical of inertial motion) are dominant. The Re number is defined as $Re = {}^{v\,L}/_{\nu}$, where $v$ is the characteristic fluid velocity, $L$ is the characteristic length of the swimmer and $\nu$ the kinematic viscosity of the fluid. In our case, given an approximated size of 10 cm and speeds of the biobots between 100-500 µm/s, we find Re numbers of the order of 1-5. In this range, both viscosity and inertia play a significant role.

**Data acquisition for the force characterization studies**

In order to evaluate the force from the cell-laden scaffold within the biobot, it was required to firstly obtain z-stack images of the posts to calculate the exact height of the tissue. The displacement of the posts upon contractions was calculated with a homemade Python script that obtained the displacement in pixels along a line perpendicular to the post border. This distance was translated into micrometers. The equation $P = \frac{3EI_z\, y(a)}{a^3}$ was used, where $P$ is the applied force, $E$ is the Young's modulus, $I_z$ is the second moment of area of the post around the z axis, $a$ is the height at which the tissue is pulling from the post and $y(a)$ is the displacement of the post at that height, was used to determine the force exerted.

**Mechanical deformation simulations**

Simulations of mechanical deformation in 3D were performed using the .stl files of the designs generated by AutoCAD. The equilibrium equations for solid mechanics given by Newton's second law were solved:

$$\rho \frac{\partial^2 \boldsymbol{u}}{\partial t^2} = \boldsymbol{\nabla} \cdot \boldsymbol{S}$$

where $\rho$ is the density of the material, $\boldsymbol{u}$ is the displacement vector and $\boldsymbol{S}$ is the second Piola-Kirchhoff stress tensor. The Young's modulus of the material was set to $E$ = 255 kPa and the poisson ratio to $\nu = 0.495$, as PDMS is nearly incompressible. Two point loads following the dynamics of a single twitch contraction, measured experimentally, were applied on both sides of the skeleton in the $x$ axis, normalized with a maximum value in the range 10-100 µN. These boundary loads model the contraction force applied by the tissue. The material was assumed to be isotropic and linear, due to the small forces and deformations that yielded von Mises stresses that were well below the yield stress of PDMS. The von Mises stress is used to predict yielding of materials under complex loading from the results obtained from uniaxial tensile test. They represent the equivalent stress across the structure and provide useful information about their distribution along the whole structure. These time-dependent equations were solved for the approximate duration of a contraction (t = 0:5 s) using the finite element method. The volume of the skeleton was meshed with tetrahedral elements and a MUMPs solver was used. To obtain the compression vs force relationship, the maximum displacement along the $x$-axis, which coincided with the time of maximum force, was computed for the left side, $x_l = \max(u_{x,l})$, and the right side, $x_r = \max(u_{x,r})$ and plotted in terms of the force. A linear least-squares fitting to the equation $x = k^{-1}F$ , allowed us to obtain the inverse of the geometrical stiffness, $k^{-1}$, for each side of the skeleton.

**Biobot kinematic model**

Hydrodynamics simulations were performed to demonstrate that symmetric spring-like skeletons should produce no net motion upon muscle contractions. Due to the high computational power of simulating the deformation of three-dimensional structures coupled with hydrodynamics, a 2D model was used instead (**Fig. 5C**). The flow fields of an incompressible liquid surrounding the symmetric design, governed by the Navier-Stokes equations, were numerically computed by FEA simulations considering the deformations that the tissue contractions apply on the skeleton. Since the deformations along the scaffold are small and can be considered linear (**Fig. 3**), the boundary conditions could be applied directly in the undeformed shape of the skeleton, thus avoiding mesh deformations and greatly reducing computational time. The contractions of the tissue were approximated by a continuous function, $g(t)$, that closely followed the shape an actual contraction profile (with a fast increase and a lower relaxation) to ensure derivability of the function. At both the left and right side of the skeleton, the deformations were defined as $d_l(t) = \Delta_l\, g(t)$ and $d_r(t) = \Delta_r\, g(t)$, respectively, where $\Delta_l$ and $\Delta_r$ were the deformations achieved at the state of maximum compression for left and right size, respectively. In this case, due to the symmetry of the design, $\Delta_l = \Delta_r$. These two values were obtained from the previous mechanical simulations of the skeleton (Figure 1.3), assuming a force of 100 µN, the same order of magnitude of the forces measured in Figure 1.4. On the rest of the biobot, namely the spring section, the deformation was approximated by a function $d(t)$, which varied linearly between $d_l(t)$ and $d_r(t)$, a characteristic that was also demonstrated by mechanical simulations. Finally, the xcomponent of the fluid flow on the biobot boundary was computed as the derivative of the deformation, $v_x(t) = \frac{d\, d(t)}{dt}$.

In more detail, as shown in **Fig. 7**, the computational domain was a square of side L from which the section of the biobot was carved out. We considered a cartesian coordinate system that fixed to the centre of the biobot. The motion of the incompressible liquid surrounding the biobot was governed by the Navier-Stokes equations:

$$\rho \frac{\partial \boldsymbol{v}}{\partial t} + \rho \boldsymbol{v} \cdot \boldsymbol{\nabla} \boldsymbol{v} = \eta \boldsymbol{\nabla}^2 \boldsymbol{v} - \boldsymbol{\nabla} p$$

$$\boldsymbol{\nabla} \cdot \boldsymbol{v} = 0$$

(1.1)

where $\rho = 1000$ kg/m$^3$ is the density of the liquid and $\eta = 10^{-3}$ Pa·s its shear viscosity. The motion of the biobot is driven by the shape deformations of the PDMS skeleton generated by the muscle tissue. These deformations generate fluid flows, which then drive the motion of the biobot. Experiments and finite elements simulations (**Fig. 3**) showed that the shape deformations were small and therefore we could apply the boundary conditions directly at its undeformed shape. This assumption greatly simplified the simulations, since mesh deformation could be avoided.

Numerical simulations of the deformation of the biobot showed that the deformation was concentrated in the spring part of the skeleton. Under the load applied by the tissue, the two curved parts on either side of the biobot were displaced as a rigid body with the deformation being concentrated in the spring. We used this finding to assume that the two curved sides were displaced as a function of time along the *x*-axis. We then assumed that the displacement varied linearly along the skeleton. The deformation on the left side was given by $d_l(t) = \Delta_l\, g(t)$ and on the right side it was given by $d_r(t) = \Delta_r\, g(t)$, where $\Delta_l$ and $\Delta_r$ are the amplitudes of the displacement of the left and of the right part of the biobot. Experimentally $\Delta_l$ and $\Delta_r$ could be different if the skeleton has a left-right asymmetric stiffness (**Fig. 3C**). The deformation on the rest of the biobot was denoted by $d(t)$and it varied linearly between $d_r(t)$ and $d_l(t)$ along the *x* coordinate, something that was also demonstrated by previous simulations (see **Fig. 7A** for a schematic representation).

The dimensionless function $g(t)$determined the time variation of the deformation. To mimic the periodic nature of the loading and relaxation of the muscle cells, we approximated the contraction by $g(t) = e^{3g^*(t)-3}$, where $g^*(t)$ is specified by the nonlinear implicit equation:

$$g^*(t) = \sin[2\pi\, f\, t + 0.8\, g^*(t)], \qquad (1.2)$$

with $f$ being the frequency of the EPS tissue stimulation. As it is shown in **Fig. 7B**, the normalized displacement prescribed by $g(t)$ closely follows that of measured experiments. The advantage of using $g(t)$ over an interpolation of the experimental data is that $g(t)$ is periodic and the average over one period of its time derivative is zero. The latter property is important to guarantee that the velocity on the boundary of the biobot caused by the deformation was zero when averaged over one period.

We assumed that the fluid velocity at the boundary of the biobot was equal to the velocity due to the deformation. Since the biobot was only deforming along the *x* coordinate and the deformations were small, we had $v_x(t) = \frac{d\, d_l(t)}{dt}$ on the left curved part, $v_x(t) = \frac{d\, d_r(t)}{dt}$ on the right curved part and $v_x(t) = \frac{d\, d(t)}{dt}$ along the rest of the skeleton. This velocity was generated by the boundary actuation of the biobot. Moreover, as we considered a reference frame that moves with the biobot, the velocity at the edges of the square domain is given by $v_x = -V_x$, where $V_x$ is the instantaneous velocity of the biobot. For simplicity, we assumed that the biobot only moves along the x axis. $V_x$ is an additional unknown that must be computed considering the balance of forces on the biobot:

$$m \frac{d}{dt} V_x = F_{h,x}, \qquad (1.3)$$

where $m$ is the mass of the biobot, which we assume has similar density as that of the liquid, and $F_{h,x}$ is the hydrodynamic force acting on it in the *x* direction. $F_{h,x}$ is computed as the integral of the stress tensor along the contour of the biobot:

$$F_{h,x} = \int_{\partial\Omega_P} \eta [\boldsymbol{\nabla v} + \boldsymbol{\nabla v}^T - p\, \boldsymbol{I}] : \boldsymbol{n}\, \boldsymbol{e}_x, \qquad (1.4)$$

where $\boldsymbol{n}$ is the vector normal to the boundary of the skeleton and pointing outwards, $\boldsymbol{e}_x$ is the unit vector along the *x* axis and the integral runs along the contour of the biobot $\partial\Omega_P$. By solving Equations 1.1 to 1.4, one finds the velocity field and the pressure field around the biobot and its velocity along the *x* axis. The equations are nonlinear because of the convective term in the Equation 1.1. We solved these equations using the finite element method. We divided the computational domain in triangular elements, with more elements near the biobot. We considered a quadratic interpolation of the velocity field and a linear interpolation for the pressure field. We used a second order implicit Runge-Kutta time integration scheme.
